# Anaplastic Thyroid Cancer Cells Upregulate Mitochondrial One-Carbon Metabolism To Meet Purine Demand, Eliciting A Critical Targetable Vulnerability

**DOI:** 10.1101/2023.04.30.538848

**Authors:** Adam J. Sugarman, Luong Do Huynh, Aidin Shabro, Antonio Di Cristofano

**Author notes:** Corresponding author: **A. Di Cristofano**, Department of Developmental and Molecular Biology, Albert Einstein College of Medicine, 1300 Morris Park Avenue, Bronx, NY 10461.

## Abstract

Anaplastic thyroid cancer (ATC) is one of the most aggressive and lethal tumor types, characterized by loss of differentiation, epithelial-to-mesenchymal transition, extremely high proliferation rate, and generalized resistance to therapy. To identify novel relevant, targetable molecular alterations, we analyzed gene expression profiles from a genetically engineered ATC mouse model and from human patient datasets, and found consistent upregulation of genes encoding enzymes involved in the one-carbon metabolic pathway, which uses serine and folates to generate both nucleotides and glycine.

Genetic and pharmacological inhibition of *SHMT2*, a key enzyme of the mitochondrial arm of the one-carbon pathway, rendered ATC cells glycine auxotroph and led to significant inhibition of cell proliferation and colony forming ability, which was primarily caused by depletion of the purine pool. Notably, these growth-suppressive effects were significantly amplified when cells were grown in the presence of physiological types and levels of folates. Genetic depletion of *SHMT2* dramatically impaired tumor growth in vivo, both in xenograft models and in an immunocompetent allograft model of ATC.

Together, these data establish the upregulation of the one-carbon metabolic pathway as a novel and targetable vulnerability of ATC cells, which can be exploited for therapeutic purposes.

## Introduction

Anaplastic thyroid cancer (ATC) is probably the most aggressive tumor type encountered in the clinic: it is typically already unresectable at presentation, highly resistant to therapy, and almost invariably lethal. ATC has a median survival of less than 9 months when patients undergo multimodal therapy, and less than 3 months with palliative care [1]. Despite being the standard of care, cytotoxic chemotherapy with taxanes, doxorubicin, or cisplatin in combination with radiation is largely ineffective in prolonging survival [2-4]. The recent FDA approval of the combination of the BRAF inhibitor dabrafenib and the MEK inhibitor trametinib for BRAF^V600E^– positive ATC represented a significant improvement in therapeutic options and patient survival[5]. Nevertheless, it benefits just a subset of patients and only for limited time, due to the rapid development of resistance [6, 7], thus underlining the urgent need for additional novel, effective, and especially rationally designed therapeutic approaches.

The one-carbon metabolism is a key process encompassing three interconnected pathways: the folate cycle, the methionine cycle, and the trans-sulfuration pathway. The folate cycle utilizes serine and dietary folate to generate glycine and tetrahydrofolate (THF)-bound one-carbon units, which are required for critical cellular needs, including de novo nucleotide biosynthesis, NADPH and glutathione production, methionine regeneration, DNA methylation and mitochondrial translation [8-10].

There are two compartmentalized arms in the folate cycle, one oxidative and mitochondrial, driven by SHMT2, MTHFD1L and MTHFD2, and one reductive and cytoplasmic, driven by SHMT1 and MTHFD1. These two arms form a generally unidirectional cycle between the two cellular compartments [11].

Notably, SHMT2 and MTHFD2 are among the most highly and consistently upregulated metabolic genes in cancer [12, 13], which underlines the cancer cells’ critical need for one-carbon-derived metabolites to fuel rapid growth and division and to withstand intracellular and extracellular oxidative stress.

The present study originated with the discovery that both mouse and human ATC display the coordinated overexpression of genes encoding key enzymes in the one-carbon pathway as well as in nucleotide biosynthesis. We have thus explored the role of the folate cycle in anaplastic thyroid cancer and demonstrate that ATC cells have an absolute and unique dependence on mitochondrial one-carbon metabolism, which appears to be essential for de novo purine biosynthesis and glycine production, but dispensable for thymidine synthesis. Furthermore, we provide in vivo evidence that targeting this pathway effectively impairs tumor growth, both in immunocompromised and in immunocompetent models of ATC.

## Materials and Methods

### Expression analyses

The datasets GSE30427 [14], GSE65144 [15] and GSE76039 [16] were analyzed using the Affymetrix Transcriptome Analysis Console (TAC) Software. The 3’scRNAseq data (GSE193581) were processed and analyzed as previously described [17].

### Cell Culture

Human ATC cell lines used in this study were maintained at 37°C with 5% CO2 in RPMI1640 (Cytiva) with 10%FBS (Biowest). For selected experiments, cells were grown in glycine-free RPMI1640 (US Biological) or folate-free RPMI1640 (Gibco). Mouse cell lines were established from ATCs developed by genetically engineered mice (manuscript in preparation) and maintained in DMEM (Cytiva) with 10%FBS. Cell line identity was validated by STS profiling (human cells) or by amplifying and sequencing genomic fragments encompassing their known mutations (mouse lines).

### shRNA-mediated gene knock-down

Cells were transduced with lentiviruses encoding shRNAs against *SHMT1* (RHS4430-101028254, RHS4430-101032591, and RHS3979-9602175), *Shmt1* (RHS4430-98486656 and RHS4430- 98714738), and *SHMT2/Shmt2* (RHS3979-9602214 and RHS3979-9602215).

ShRNA RHS3979-9602214 was also cloned into the Tet-pLKO-puro (Addgene plasmid #21915). Lentiviral constructs were packaged in HEK293 cells transfected using Lipofectamine 2000 (Invitrogen). Viral supernatant was collected 48-72 hours after transfection, combined, and filtered using 0.45 μm Nalgene SFCA syringe filters (Thermo Scientific). Target cells were plated 24 hours before infection and then exposed to viral supernatant supplemented with 20 μg/ml polybrene (Santa Cruz). Mouse cell lines were then selected for at least 72 hours with 4 μg/ml puromycin (Corning), whereas a concentration of 2 μg/ml was used for human cell lines.

Transduced cell lines were pretreated with 1 µg/ml doxycycline hyclate (Sigma Aldrich) for 48 hours prior to plating for growth curve experiments. In all experiments performed on transduced cell lines, doxycycline hyclate was replaced every 48 hrs.

### Drugs and chemicals

The following chemical compound were used: SHIN1 (Aobius), DS18561882 (Medchemexpress), Hypoxanthine (TCI America), Thymidine and Glycine (Sigma Aldrich), Sodium formate (Thermo Scientific), ethyl ester GSH, methotrexate, pemetrexed, folic acid and 5-methyl-THF (Cayman Chemical).

### Viability assays

Cells were plated in 96-well plates in medium with either FBS or dialuzed FBS (Biowest). Treatments were added 24 hours after plating. Alamar Blue was directly added to the culture medium of treated and control cells after 72 hours of treatment. Fluorescence was measured using a plate reader (excitation 530nm, emission 590nm). Statistical analysis and EC50 value calculation were obtained using GraphPad Prism (GraphPad Software).

For cell counts, cells were collected at the end of treatment and counted using a Z2 Coulter particle counter (Beckman).

Synergy analysis was performed using the ZIP score method in Synergy Finder ((https://synergyfinder.fimm.fi) [18].

### Colony formation assay

250 cells were plated in each well of a 12-well plate. Treatments were added 24 hours after plating. Medium was replaced every 72 hours. Colonies were fixed with 10% neutral buffered formalin and stained with 0.01% crystal violet (Sigma-Aldrich).

### Cell Cycle analysis

For cell cycle analysis, cell lines were harvested by trypsin treatment and fixed in 75% ethanol in ice for 30 min. After treatment with RNase (Roche) for 5 min at room temperature, cells were stained with propidium iodide (BioSure) overnight, and DNA content was measured using a BD FACSCantoTM II system (BD Biosciences).

### Western blot

Cells were homogenized on ice in radioimmunoprecipitation assay (RIPA) buffer supplemented with Halt Protease and Phosphatase Inhibitor cocktail (ThermoFisher Scientific). Protein concentration was determined using the Pierce BCA protein Kit (ThermoFisher Scientific). Western blot analysis was conducted using 50 µg of proteins on ExpressPlus precast gels (Genescript). Proteins were blotted onto polyvinylidene difluoride membranes (Millipore). The membranes were probed with the indicated antibodies (all from Cell Signaling). All the primary antibodies were used at a dilution of 1:1000 in 5% BSA in TBS-T. Signals were detected with HRP-conjugated secondary antibodies (ThermoFisher Scientific) and the chemiluminescence substrate Luminata Crescendo (EMD Millipore).

### In vivo studies

6-8 week-old NOD-*scid IL2r^null^* (NSG) mice (JAX) were injected subcutaneously with 5×10^6^ cells. Where needed, Doxycycline hyclate was dissolved in drinking water (2mg/ml) and replaced every three days. Tumor volume was calculated from two-dimensional measurements using the equation: tumor volume = (length × width^2^) × 0.5, every three days. Tumor weight was measured at the end of the experiment. Data were plotted and analyzed using GraphPad Prism.

All the animal studies were approved by the Einstein Institutional Animal Care and Use Committee.

## Results

Analysis of an RNA expression dataset (GSE30427) generated from ATCs developed by thyroid-specific (*Pten*,*Tp53*)^-/-^ mice [14] revealed the coordinated overexpression of most of the genes that encode enzymes involved in the serine biosynthesis, one-carbon, and de novo purine biosynthesis metabolic pathways (Fig. 1A,D and Table 1). Overexpression of *SHMT1*, *SHMT2*, and *MTHFD2*, three key players in the one-carbon pathway for which small molecule inhibitors have recently been developed [19, 20], was confirmed at the protein level using extracts from wild type thyroids and from tumors developed by three different ATC mouse models, [*Pten*,*Tp53*]^-/-^ [14], *Nras*^Q61R^, *Pten^-/-^*,*Tp53^-/-^*, and *Nras*^Q61R^, *PIK3CA^H1047R^*,*Tp53^-/-^*(manuscript in preparation) (Fig. 1B).

**Figure 1.**
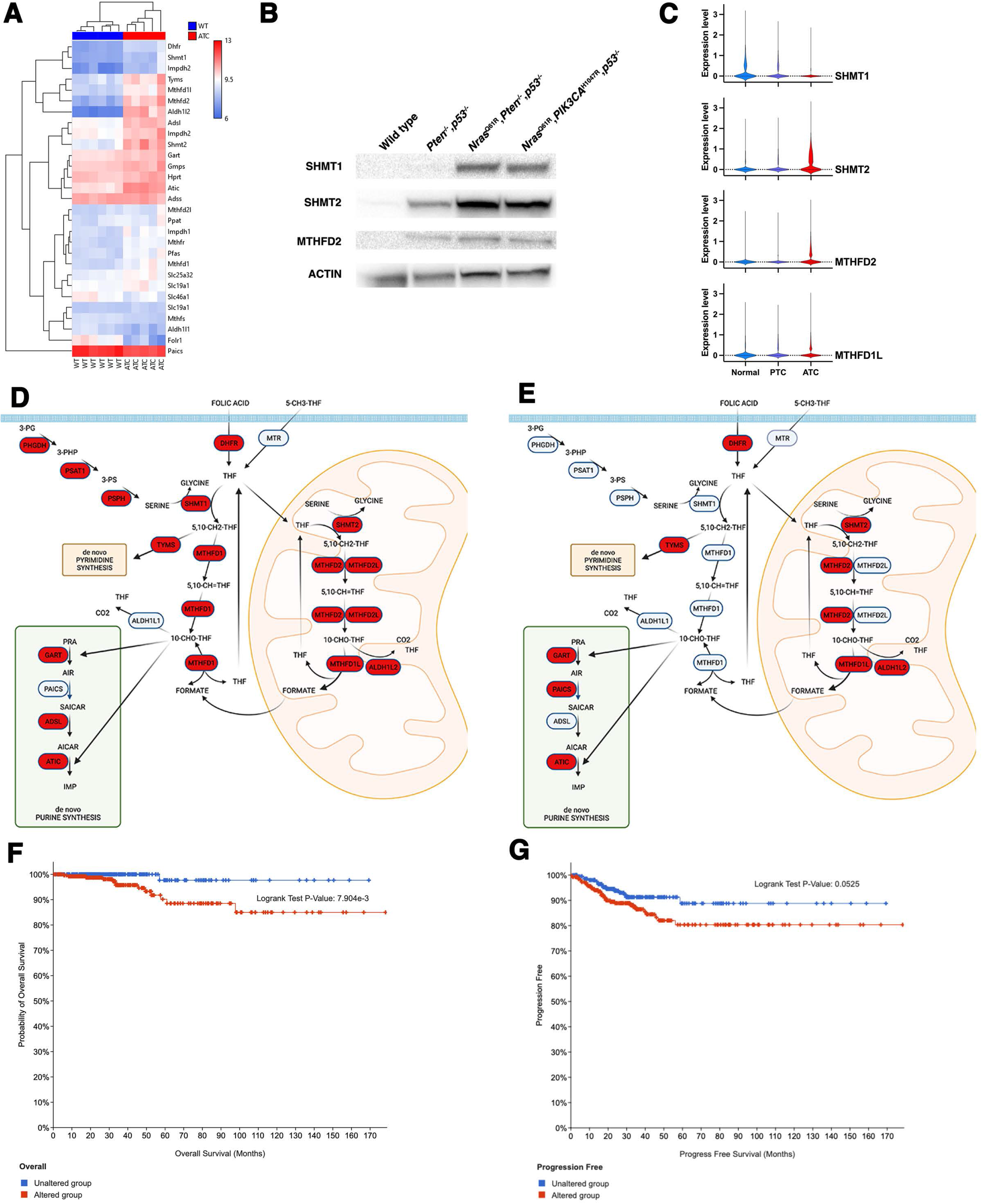
One-carbon metabolism genes are overexpressed in anaplastic thyroid cancer. A) Hierarchical clustering analysis of gene expression for one-carbon, purine synthesis, and thymidine synthesis genes from wild type thyroids and ATCs developed by [*Pten,Tp53*]^-/-^ mice. B) Western blot analysis of protein extracts from wild type thyroids and ATCs developed by mice of the indicated genotypes. C) Violin plot analysis of scRNAseq data from normal thyroids, PTC and ATC. D) Graphic representation of the overexpression (in red) of serine synthesis, one-carbon, purine synthesis, and thymidine synthesis genes in mouse (left) and human (right) ATCs, as detected by microarray expression analysis. F,G) Overall and progression-free survival of PTC patients overexpressing *SHMT2* and *MTHFD2* (from TCGA data).

**Table 1.**

| **Gene name** | **Mm (GSE30427)** | **Hs1 (GSE65144)** | **Hs2 (GSE76039)** |
| --- | --- | --- | --- |
| One-carbon metabolism |  |  |  |
| ***DHFR*** | **1.076** | **0.748** | **1.465** |
| *SHMT1* | 0.767 | -1.356 | -1.470 |
| ***SHMT2*** | **1.684** | **1.795** | **1.714** |
| *MTHFD1* | 0.79 | 1.029 | -1.362 |
| ***MTHFD1L*** | **1.359** | **1.465** | **2.144** |
| ***MTHFD2*** | **2.625** | **2.195** | **2.751** |
| *MTHFD2L* | 0.459 | 0.322 | -1.561 |
| *ALDH1L1* | -0.718 | 0.310 | nc |
| ***ALDH1L2*** | **3.86** | **1.722** | **0.465** |
| Nucleotide synthesis |  |  |  |
| ***TYMS*** | **1.587** | **2.888** | **3.966** |
| ***GART*** | **0.363** | **0.918** | **0.986** |
| *PAICS* | -0.509 | 1.007 | 1.541 |
| *ADSL* | 0.963 | 0.465 | nc |
| ***ATIC*** | **1.01** | **0.595** | **0.978** |

Log<sub>2</sub> fold change vs. normal thyroid. In bold, genes overexpressed in all datasets

We cross-referenced the list of serine/one-carbon/purine metabolism genes deregulated in mouse ATCs with two datasets derived from human controls and ATC patients (GSE65144 and GSE76039) and found that human tumors selectively upregulate genes involved in the mitochondrial arm of the one-carbon pathway and genes in the de novo purine biosynthesis pathway (Table 1). Finally, we used a recently published scRNAseq dataset from normal thyroids, papillary thyroid cancer (PTC), and ATC patients [Lu, 2023] to further validate this pattern of overexpression: in fact, SHMT2 MTHFD2 and MTHFD1L were selectively upregulated in ATC (Fig. 1C).

Increased expression of SHMT2 and MTHFD2, but not SHMT1, was also confirmed at the protein level using several human ATC cell lines, compared to the immortalized NTHY-Ori human thyroid line (Suppl. Fig. 1).

These results indicate that both mouse and human ATC cells upregulate the mitochondrial component of the one-carbon pathway as well as several genes involved in the de novo purine synthesis pathway. In addition, mouse cells also upregulate the cytoplasmic component of the one-carbon pathway and the serine biosynthesis pathway (Fig. 1D).

To gain clinical insight on the significance of this finding, in the absence of an equivalent resource for ATCs, we analyzed the TCGA database, which contains data from PTC patients, and found that coordinated overexpression of *MTHFD2* and *SHMT2* is significantly associated with reduced overall survival and progression-free survival, further underlining the clinical relevance of these molecular alterations (Fig. 1E,F).

In order to characterize the biological consequences of the overexpression of one-carbon pathway genes, we generated mouse and human ATC cell lines expressing control shRNAs or shRNAs targeting *Shmt1*/*SHMT1* and/or *Shmt2*/*SHMT2*. Strikingly, while we could readily generate mouse cells constitutively expressing these constructs, human cells did not survive constitutive *SHMT2* targeting, and we had to use a doxycycline-inducible vector for these shRNAs. This species difference likely reflects the fact that human cells do not overexpress *SHMT1*, and thus cannot compensate the effects of *SHMT2* depletion. More importantly, the inability to survive in the prolonged absence of *SHMT2* indicate that human ATC cells are uniquely addicted to the one-carbon metabolic pathway.

When the engineered ATC cells were grown in the presence of normal fetal bovine serum (FBS), which contains several of the small metabolites whose levels are affected by targeting the one-carbon pathway (in particular nucleotides), depletion of *Shmt2*/*SHMT2* significantly reduced the proliferation of both mouse and human cells. However, while depletion of *Shmt1* slightly reduced the proliferation of mouse cells, targeting *SHMT1* did not alter the proliferation of human ATC cells, strongly suggesting a direct correlation between isoform overexpression and its requirement (Fig 2A,B and Suppl. Fig. 2). Interestingly, co-targeting both genes did not result in further decreased proliferation (Fig. 2A,B), even in the mouse cells, suggesting that *Shmt2*/*SHMT2* function is the major determinant of ATC cells proliferative activity.

**Figure 2.**
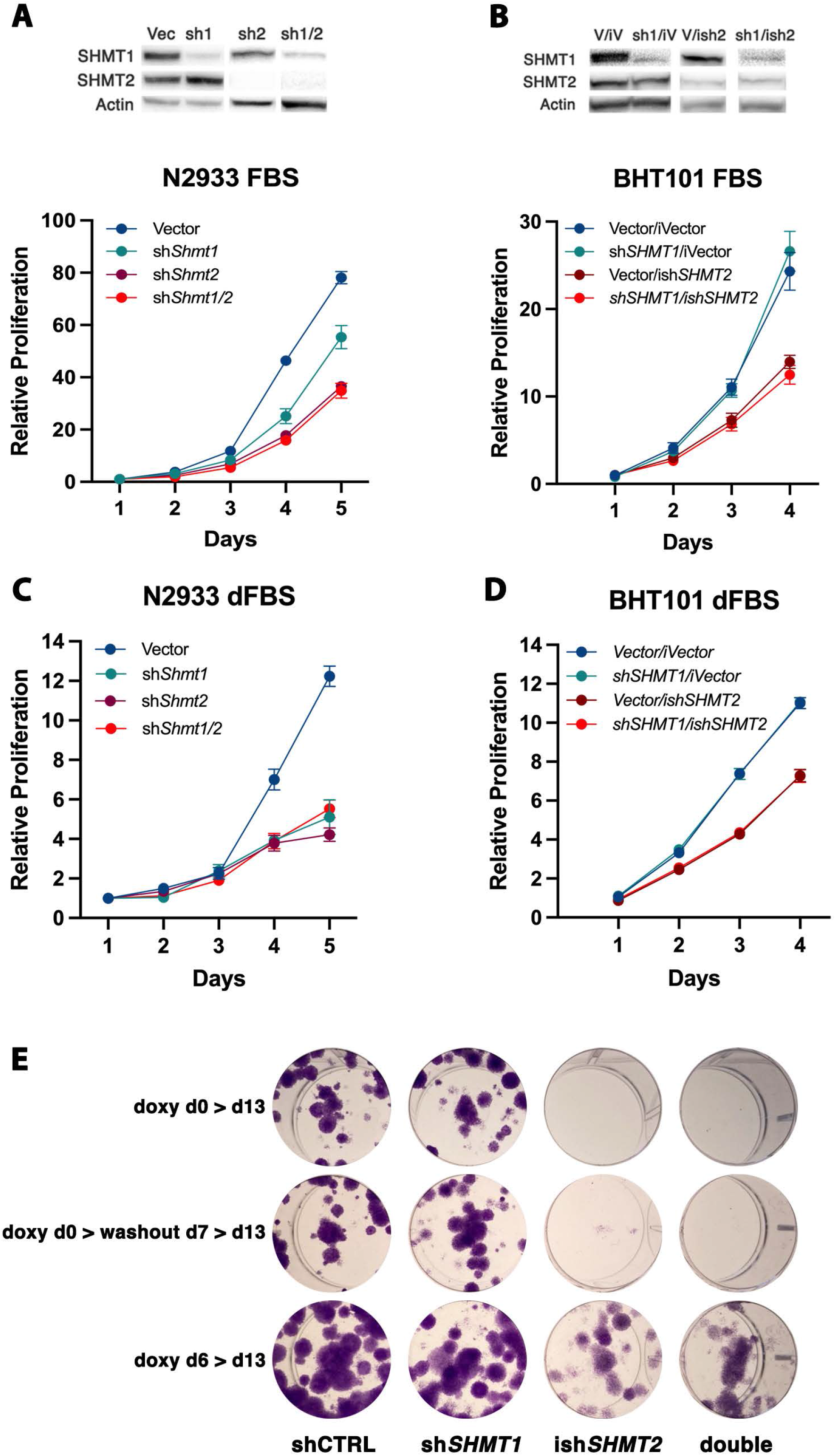
Genetic downregulation of SHMT2 impairs proliferation of ATC cells. A,B) Representative analysis of cell proliferation of mouse (A) and human (B) ATC cell lines transduced with non-targeting shRNA, sh*SHMT1*, sh*SHMT2*, sh*SHMT1/2*. For the human cells, doxycycline-inducible sh*SHMT2* was used. Similar results were obtained with a second set of shRNAs (not shown). C,D) Same experiment as A-B, but using RPMI1640 with dialyzed FBS. E) Colony forming assay of OCUT2 cells expressing the indicated shRNAs. Expression of sh*SHMT2* was induced at day 0 (top and middle) or at day 6. In the middle panel, doxycycline was removed at day 7 to allow re-expression of sh*SHMT2*.

When the mouse ATC cells were grown in the presence of dialyzed FBS, in which small metabolites have been removed, depletion of *Shmt1* reduced mouse ATC cell proliferation to the same extent as depletion of *Shmt2*, suggesting that, in the absence of an extracellular supply of metabolites, high expression of both isoforms is required to support cell proliferation (Fig. 2C). Conversely, human ATC cells still were unaffected by *SHMT1* depletion, in line with the absence of alterations in the expression level of this gene in human samples (Fig. 2D).

To define the long term impact of genetic inhibition of the one-carbon pathway, we performed colony-forming assays using two human ATC cell lines, BHT101 and OCUT2, either in the presence of regular FBS or of dialyzed serum (Fig. 2E and Suppl. Fig. 3). In line with the results of the short-term cell proliferation assays, shRNA-mediated depletion of *SHMT1* did not alter the ability of ATC cells to form colonies. Conversely, depletion of *SHMT2* dramatically inhibited colony formation (Fig. 2E and Suppl. Fig. 3, top). As expected, this inhibitory effect was significantly amplified by the absence of rescuing metabolites in the dialyzed serum.

To test whether the growth arrest induced by *SHMT2* depletion is reversible, we washed out doxycycline after one week and let the cells grow for an additional week. Strikingly, re-expression of *SHMT2* did not result in restored cell proliferation, strongly suggesting that one-carbon pathway inhibition exerts long-lasting inhibitory effects on ATC cell proliferation (Fig. 2E and Suppl. Fig. 3, middle).

Finally, we tested whether depleting *SHMT2* after colony formation was able to prevent further colony growth. To this end, doxycycline was added at day six, after the appearance of well-established colonies. Once again, *SHMT2* depletion resulted in significantly reduced colony size, compared to controls, indicating that one-carbon pathway inhibition may be effective on already established tumors (Fig. 2E and Suppl. Fig. 3, bottom).

Over the past few years, novel selective inhibitors of enzymes of the one-carbon pathway have been developed. In particular, SHIN1 has been synthesized as an inhibitor of SHMT1/2 [19], and DS18561882 as an inhibitor of MTHFD2 [20]. These compounds have recently been shown to reduce cell proliferation in lymphoma [19], rhabdomyosarcoma [21], bladder [22] and lung cancer [23, 24] cell lines.

We tested the effect of these inhibitors on a large panel of mouse and human ATC cell lines encompassing the entire spectrum of driver mutations associated with ATC development.

SHIN1 was extremely effective in inhibiting the proliferation of both mouse and human ATC cells, with EC50s in the low micromolar range (Fig. 3A,B). Interestingly, cell lines carrying RAS activating mutations (CAL62, ACT1, Hth7, C643) were particularly sensitive to the inhibitory effect of SHIN1 (Fig. 3B, blue lines).

**Figure 3.**
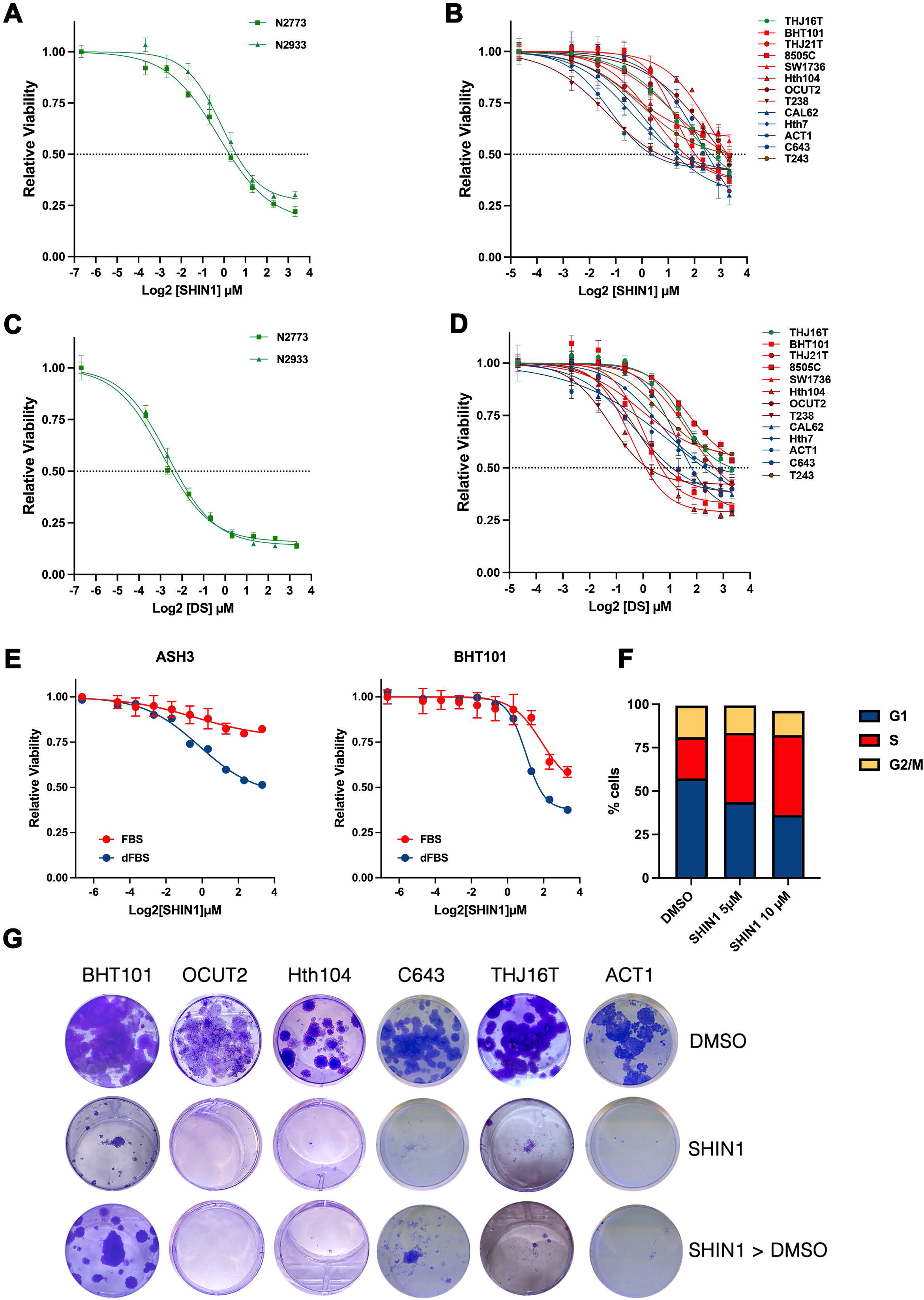
Small molecule inhibition of the one-carbon metabolism impairs proliferation of ATC cells. A-D), Viability of mouse (A) and human (B) ATC cell lines exposed to the SHMT1/2 inhibitor SHIN1 (A,B) and the MTHFD2 inhibitor DS18561882 (C,D). In B and D, cell lines are color coded based on their driver genetic alterations: mutant BRAF (red), mutant RAS (blue), mutant PI3K (green), mutant BRAF and PI3K (maroon), mutant BRAF and RB1 (brown). E) Effect of dialyzed FBS on the response of ATC cells to SHMT1/2 inhibition. F) Analysis of cell cycle distribution of BHT101 cells treated with SHIN1 for 72h. G) Representative colony forming assays of six human ATC cell lines treated for two weeks with 10µM SHIN1. In the last row, SHIN1 was removed after one week.

Both mouse and human ATC cell lines were also very sensitive to the growth-inhibitory effect of DS18561882, with EC50s in the high nanomolar-low micromolar range (Fig. 3C,D). However, contrary to SHIN1, we could not find any significant correlation between inhibitor sensitivity and driver genetic alteration.

Combined inhibition of SHMT1/2 and MTHFD2 resulted in moderate synergy, as calculated using the ZIP model, in line with the notion that co-targeting enzymes in a linear pathway does not result in a dramatic improvement over single target inhibition (Suppl. Fig. 4).

The effect of pharmacological inhibition of the one-carbon pathway was significantly amplified when the experiments were conducted in dialyzed FBS, once again suggesting that the observed growth inhibitory effect is linked to the production and availability of small metabolites (Fig. 3E).

To identify the mechanisms leading to growth suppression upon inhibition of the one-carbon metabolic pathway, we measured the effect of pharmacological SHMT1/2 inhibition on both apoptosis and cell cycle distribution in the BHT101 cell line. While SHIN1 treatment did not result in any induction of apoptosis (not shown), we observed a dose-dependent accumulation of cells in S phase, suggesting that SHIN1 impairs the ability of ATC cells to complete DNA replication (Fig. 3F).

In order to assess the effect of long-term SHMT1/2 inhibition, we performed colony-forming assays using a panel of ATC lines encompassing all the oncogenic driver combinations commonly observed in human patients and treated them with 10µM SHIN1, a concentration roughly corresponding to the 72h EC70. Two weeks after continuous drug exposure, colony formation was completely suppressed by SHMT1/2 inhibition (Fig. 3G, middle vs. top). To assess the reversibility of the growth suppression consequent SHMT1/2 inhibition, we treated the cells with SHIN1 for one week and then removed the compound for another week. Strikingly, five out of six cell lines completely failed to resume proliferation (Fig. 3G), while BHT101 cells, which express the highest levels of SHMT2 (Fig 1C), showed a modest resumption of proliferation. These data confirm the genetic approaches described in Fig.2 and strongly suggest that, in ATC cells, one-carbon pathway inhibition leads to a prolonged state of growth suppression.

Further analysis of the ATC expression dataset GSE76039, in which the driver mutations are known for each tumor, showed that the expression of both *SHMT2* and *MTHFD2* is significantly higher in RAS-driven than in BRAF-driven thyroid anaplastic tumors (Fig. 4A). To assess the significance of this association, we performed shRNA-mediated depletion of *NRAS* in the ACT-1 ATC cell line (carrying a Q61K NRAS mutation) and found that it significantly reduced its sensitivity to SHIN1 treatment (Fig. 4B). Furthermore, inhibition of MAPK signaling using the MEK inhibitor, trametinib, exerted the same desensitizing effect on mutant RAS-driven cell lines (Fig. 4C) but not on mutant BRAF-driven ones (data not shown). Furthermore, the sensitivity of ATC cell lines to SHMT1/2 inhibition was found to correlate with their RAS score, which uses a 71-gene expression signature to quantify the extent to which the gene expression profile of a given tumor resembles either the *BRAF*^V600E^ or *RAS*-mutant profiles [25] (Fig. 4D). These data strongly suggest the presence of a functional association between RAS activation and SHMT2 overexpression and requirement. Finally, we also found a significant inverse correlation between the sensitivity of ATC cell lines to SHMT1/2 inhibition and their Thyroid Differentiation Score (TDS) [25], with more dedifferentiated cell lines being more sensitive to SHIN1 treatment (Fig. 4E).

**Figure 4.**
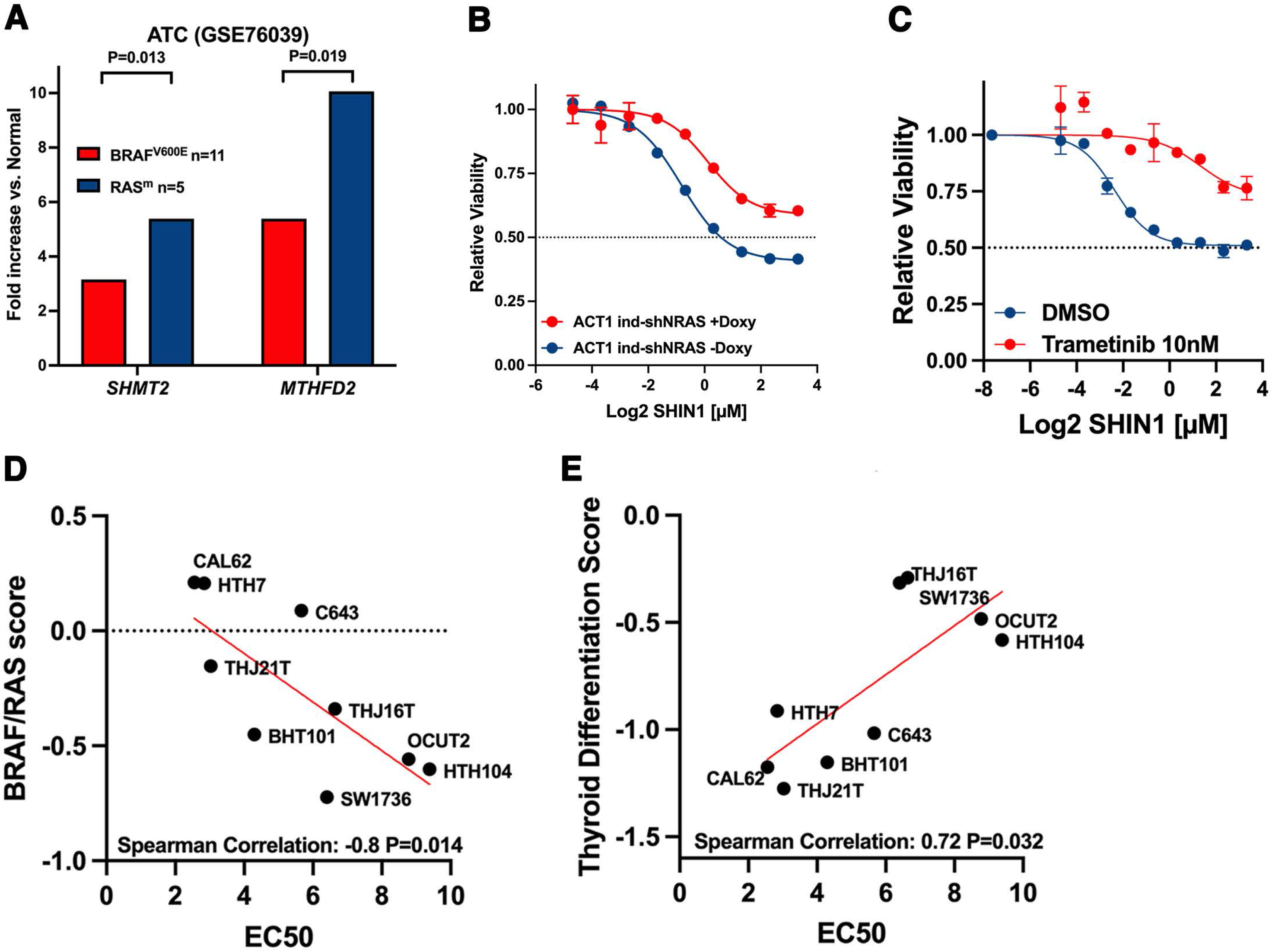
RAS mutant cells express higher levels of SHMT2 and MTHFD2 than BRAF mutant cells. A) Average increase of *SHMT2* and *MTHFD2* expression in human ATCs by driver mutation. B) Effect of mutant NRAS downregulation in ACT1 cells on sensitivity to SHIN1. C) Effect of MEK inhibition in ACT1 cells on sensitivity to SHIN1. D) Correlation between ATC cell line BRAF/RAS score (higher is more RAS-like) and sensitivity to SHIN1. E) Correlation between ATC cell line thyroid differentiation score (lower is less differentiated) and sensitivity to SHIN1.

Inhibition of the one-carbon metabolic pathway leads to two critical deficiencies: depletion of formate, a key intermediate produced by the mitochondrial arm of the pathway, which carries the one-carbon unit into the cytoplasm to fuel de novo nucleotide synthesis, and inhibition of the major pathway for intracellular glycine synthesis (Fig. 1D).

To define the relative impact of one-carbon metabolism inhibition on these pathways in ATC cells, we first tested the rescuing effect of exogenous formate on the growth-suppression consequent pharmacological inhibitors of this pathway in BHT101 and C643 cells. Supplementation of the growth medium with 1mM formate completely restored the proliferative ability of ATC cells despite the presence of SHMT1/2 and MTHFD2 inhibitors (Fig. 5A, Suppl. Fig. 5A). To distinguish between the effect of one-carbon metabolism inhibition on purine synthesis and that on thymidine synthesis, we tested the rescuing effect of exogenous hypoxanthine, which feeds the purine salvage pathway, and that of thymidine. Strikingly, only hypoxanthine rescued the proliferative block induced by both SHIN1 and DS18561882, strongly suggesting that, in ATC cells, purine depletion is the critical consequence of one-carbon metabolism inhibition (Fig. 5B, Suppl. Fig. 5B).

**Figure 5.**
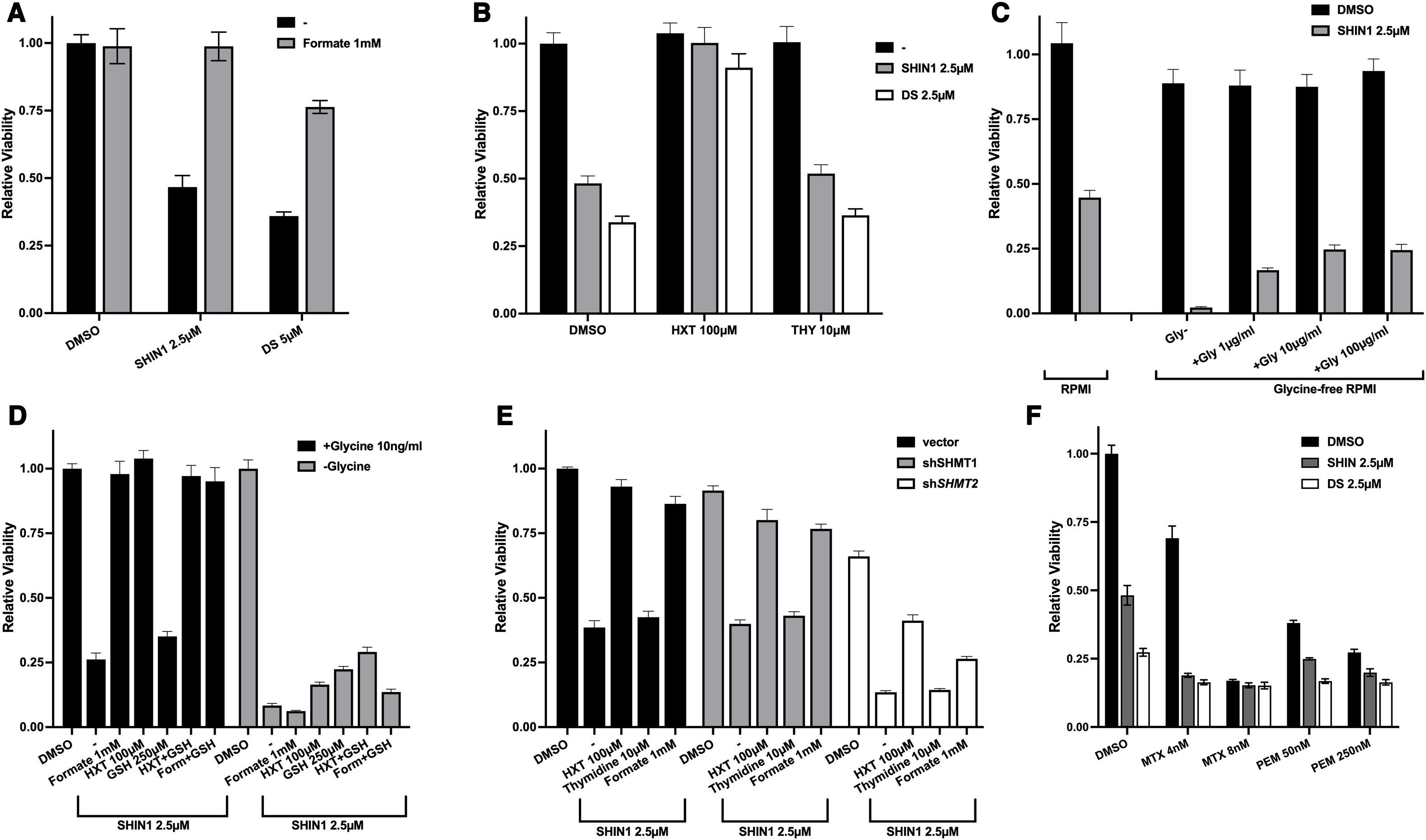
Functional analysis of one-carbon pathway inhibition in BHT101 cells. A) Rescuing effect of exogenous formate on the growth inhibition induced by one-carbon pathway inhibitors. B) Exogenous hypoxanthine, but not thymidine, rescues the growth inhibition induced by one-carbon pathway inhibitors. C) Lethal effect of exogenous glycine deprivation combined with SHIN1 and rescuing effect of exogenous glycine added to the medium. D) Systematic analysis of the extent of viability rescue in the presence of exogenous formate, hypoxanthine, and glutathione (GSH) in the presence and absence of extracellular glycine. E) Analysis of the ability of exogenous formate, hypoxanthine, and thymidine to rescue the inhibitory effect of SHIN1 in cells depleted of *SHMT1* or *SHMT2*. F) Effect of combining one-carbon pathway inhibitors with different concentrations of methotrexate (MTX) and pemetrexed (PEM).

When we inhibited SHMT1/2 in glycine-free medium, viability of ATC cells dropped below the assay detection level and was almost completely rescued by exogenous glycine supplementation (Fig. 5C). Notably, direct cell counting experiments demonstrated that, in the absence of glycine, ATC cells undergo cell death when one-carbon metabolism is inhibited (Suppl. Fig. 5C). These data clearly demonstrate that ATC cells in which SHMT-mediated glycine synthesis is inhibited become auxotroph for this amino acid.

Besides its role in nucleotide production, cells need glycine to support glutathione synthesis to fight the various sources of oxidative stress they are exposed to. We found that exogenous glutathione added to the culture media in the presence of glycine was unable to rescue the proliferative block of ATC cells. However, in the absence of glycine, glutathione and hypoxanthine were able to partially rescue viability in an additive manner (Fig. 5D, Suppl. Fig. 5D). Thus, while extracellular glycine, in the absence of endogenous glycine synthesis, appears to be sufficient to sustain sufficient glutathione synthesis, under complete glycine deprivation conditions ATC cell death seems to be partially caused by glutathione depletion. Interestingly, the incomplete viability rescue observed when supplementing glutathione and hypoxanthine suggests the existence of additional critical roles for glycine.

To further validate the relative role of SHMT1 and SHMT2, we performed the nucleotide rescue experiments in the BHT101 line expressing shRNAs for these two genes. As expected, sh*SHMT1*- expressing cells behaved exactly like the control cells, supporting the notion that SHMT1 does not play a key role in ATC one-carbon metabolism. Conversely, sh*SHMT2*-expressing cells were more sensitive to SHIN1 treatment. Notably, both hypoxanthine and formate were unable to fully rescue the proliferative defect of these cells, suggesting that this more profound SHMT2 inhibition (shRNA plus inhibitor) might be uncovering additional SHMT2 functions not related to purine biosynthesis (Fig. 5E).

Finally, we tested the effect of combining SHIN1 with two folate analogues, methotrexate, which inhibits DHFR and thus acts upstream of the one-carbon pathway, and pemetrexed, which inhibits thymidylate synthase and, secondarily, phosphoribosylglycinamide formyltransferase (GART). As expected for a linear pathway, low doses of methotrexate synergized with SHIN1, while the response to higher doses, which lead to complete pathway inhibition, was not modified by SHMT1/2 inhibition. On the contrary, pemetrexed had only a mildly additive effect, confirming the notion that thymidine synthesis is not a critical conduit of SHMT1/2 inhibition (Fig. 5F).

Commonly used cell culture media contain very high levels of folic acid as a precursor of THF. These extreme, non-physiological conditions are likely to affect the response of cultured cells to one-carbon pathway inhibitors. To test the effect of SHIN1 on ATC cells in more clinically relevant conditions, we grew them in folate-free medium for four days to reduce the endogenous levels of folates, and then generated dose-response curves either in regular RPMI (2.27µM folic acid) or in folate-free RPMI supplemented with 50nM 5-methyl-THF, the primary source of dietary folate. Strikingly, all four ATC cell lines tested displayed significantly increased sensitivity to low levels of SHIN1, strongly suggesting that, in physiological conditions, ATC cells are even more sensitive to inhibition of one-carbon metabolism (Fig. 6A).

**Figure 6.**
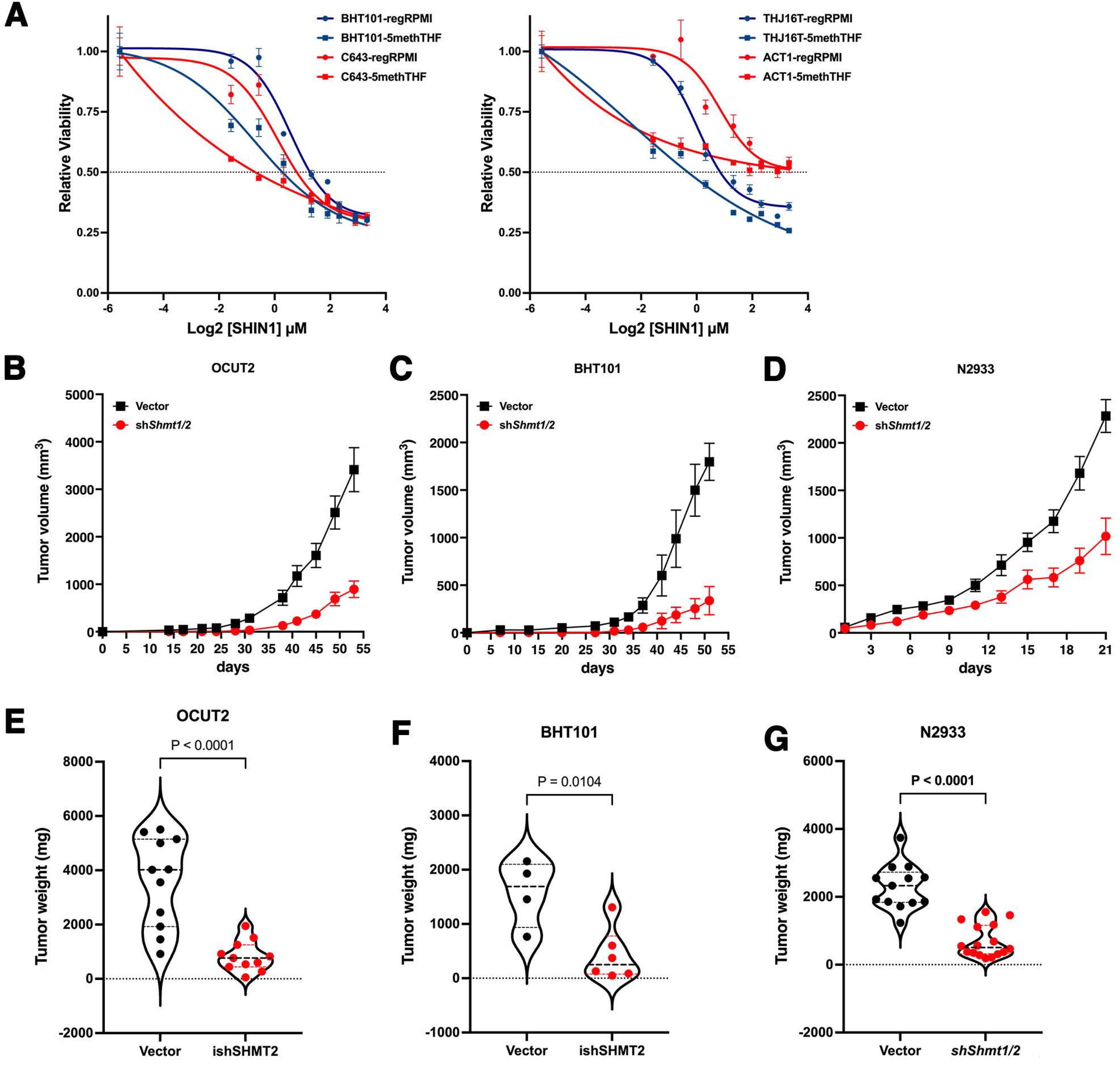
Effect of physiological folates on the efficacy of one-carbon metabolism inhibitors and of *SHMT1/2* depletion on tumor growth in vivo. A) Low concentrations of SHIN1 are strikingly more effective when ATC cells are grown in the presence of 50nM 5-methyl-THF instead of 2.27µM folic acid. B-G) shRNA-mediated depletion of *SHMT1* and *SHMT2* severely impairs in vivo tumor growth in OCUT2 (B,E) and BHT101 (C,F) xenografts, as well as in immunocompetent N2933 (D,G) allografts, resulting in significantly smaller tumors.

While SHIN1 is an excellent tool compound for in vitro studies, it has a very poor bioavailability [19]. Thus, to assess in vivo the effect of inhibiting one-carbon metabolism in ATC cells, we injected two different cell lines, OCUT2 and BHT101, expressing either a control shRNA or sh*SHMT1* (constitutive) plus sh*SHMT2* (doxycycline-inducible), into NSG mice. Mice were injected on one side with the control cells and on the other side with the SHMT1/2-depleted cells. Both SHMT-depleted cell lines showed a dramatic reduction in tumor growth rate (Fig. 6B, C) as well as in tumor weight at the end of the experiment (Fig. 6E, F), thus providing further proof that inhibition of one-carbon metabolism could be a viable strategy for ATC patients. To further corroborate these data in an immunocompetent model, we also generated allografts using mouse ATC cells expressing either a control construct or shRNAs targeting *Shmt1* and *Shmt2*. Once again, the SHMT-depleted cells were drastically impaired in their ability to grow in vivo (Fig. 6D) and developed significantly smaller tumors (Fig. 6G).

## Discussion

Metabolic alterations taking place in cancer cells to cope with their increased proliferative activity as well as with the unique microenvironment they inhabit often represent targetable liabilities that can be harnessed for therapeutic purposes.

Our data clearly demonstrate that increased expression of one-carbon metabolism genes is a major driver of thyroid anaplastic cancer cells proliferation, and that it represents a remarkable addiction, probably unique to this tumor type. In fact, to our knowledge, ATC cells are the only cancer cells studied so far that cannot withstand long-term inactivation of one-carbon metabolism. This notion is first of all supported by the inability to develop cell lines stably expressing shSHMT2 constructs, in contrast with what has been described for T-ALL [26], renal cell carcinoma [27], glioblastoma [28], and breast cancer [29]. Second, our data provide evidence that the effect of genetic or chemical inhibition of one-carbon metabolism in ATC cells has strong cytostatic effects that persist for at least for one week even when the inhibition is removed. Whether these cells enter senescence, or a prolonged but reversible state of quiescence remains to be addressed in future work.

It is not clear, at this stage, why mouse cells upregulate both the cytoplasmic and the mitochondrial arms of the folate cycle, while human cells only increase the mitochondrial one. It is possible that this difference is correlated with the much higher proliferation rate of mouse ATC cells compared to the human ones, requiring additional metabolic adaptations to sustain the replicative activity.

Our finding that RAS mutant ATC cells express higher levels of *SHMT2* and *MTHFD2* than BRAF mutant cells, coupled with the observed correlation between the presence of mutant RAS and the sensitivity to one-carbon pathway inhibition, confirms and extends previous data showing that RAS-transformed fibroblasts increase the expression of one-carbon metabolism enzymes and production of formate [30]. It is possible that mutant RAS signals to stimulate the coordinate transcription of these genes, leading to increased metabolic activity and addiction, and thus sensitivity to pathway inhibition. Future studies will need to address the mechanistic link between RAS and one-carbon metabolism dependency, an area of study that might lead to a rational stratification of thyroid cancer patients by their predicted sensitivity to one-carbon metabolism inhibitors.

We have shown that hypoxanthine, but not thymidine, rescues the proliferation block associated with inhibition of one-carbon metabolism. While in agreement with other studies, where purine synthesis is the specific essential conduit of SHMT1/2 inhibition [19], these data are in stark contrast with some other models, where it is impaired thymidine synthesis that appears to be the critical downstream consequence of one carbon metabolism inhibition [31]. Whether these differences are due to the specific wiring of the cell types analyzed, or to the experimental approaches utilized, remains to be elucidated.

Inhibition of one-carbon metabolism leads to glycine auxotrophy and rapid cell death when cells are deprived of extracellular sources of glycine. It is not known which amino acid transporter(s) is responsible for glycine entry in ATC cells. It is tempting to speculate that identification of a relatively cell type-specific transporter would open the possibility of generating a synthetic lethal therapeutic approach.

While most of the experiments described in our study, and basically all those available in the literature, have been performed using cell culture media containing extremely high levels of folic acid, which is not the main source of dietary folates, and certainly not present in cells at such high concentrations, our data show that using medium supplemented with physiological concentrations of 5-methyl-THF leads to an even higher sensitivity of ATC cells to one-carbon pathway inhibition. This notion suggests that the in vivo efficacy of one-carbon pathway inhibitors would be higher than what we can infer from current in vitro data.

SHIN1, the best characterized and most selective SHMT1/2 inhibitor available, is not suited for in vivo studies due to its rapid clearance. A more bioavailable derivative, SHIN2, has recently been described [32]. While this compound shows limited and reversible hematological toxicity and promising activity in T-ALL models, it is extremely expensive, which has prevented us from testing it in our models.

Thus, as a proof of principle, we have used genetic approaches to inhibit one-carbon metabolism in vivo. Our data from two immunocompromised xenograft models and one immunocompetent allograft model consistently suggest that SHMT inhibition is a viable and promising strategy to approach ATC. While inhibition of one-carbon metabolism in ATC cells appears to be cytostatic, the prolonged growth arrest observed in cell culture suggests the possibility of extended tumor control in vivo, a therapeutic goal of utmost importance in ATC management. In addition, these approaches represent a cornerstone for the development of rationally designed synthetic lethal strategies harnessing the liabilities exposed by one-carbon metabolism impairment.

## Acknowledgements

Research reported in this publication was supported by NIH grant CA128943 to ADC and by a Ruth L. Kirschstein T32 Training Grant of Surgeons for the Study of the Tumor Microenvironment (CA200561). We acknowledge the Animal Housing Core Facility of Albert Einstein College of Medicine, which is partially supported by the NIH Cancer Center Support Grant to the Albert Einstein Cancer Center (P30CA013330).

We thank Dr. David Goldman for providing invaluable input and advice, as well as Dr. Ruli Gao and Dr. Lina Lu for providing the scRNAseq data.

## Figure legends

**Suppl. Fig. 1.**
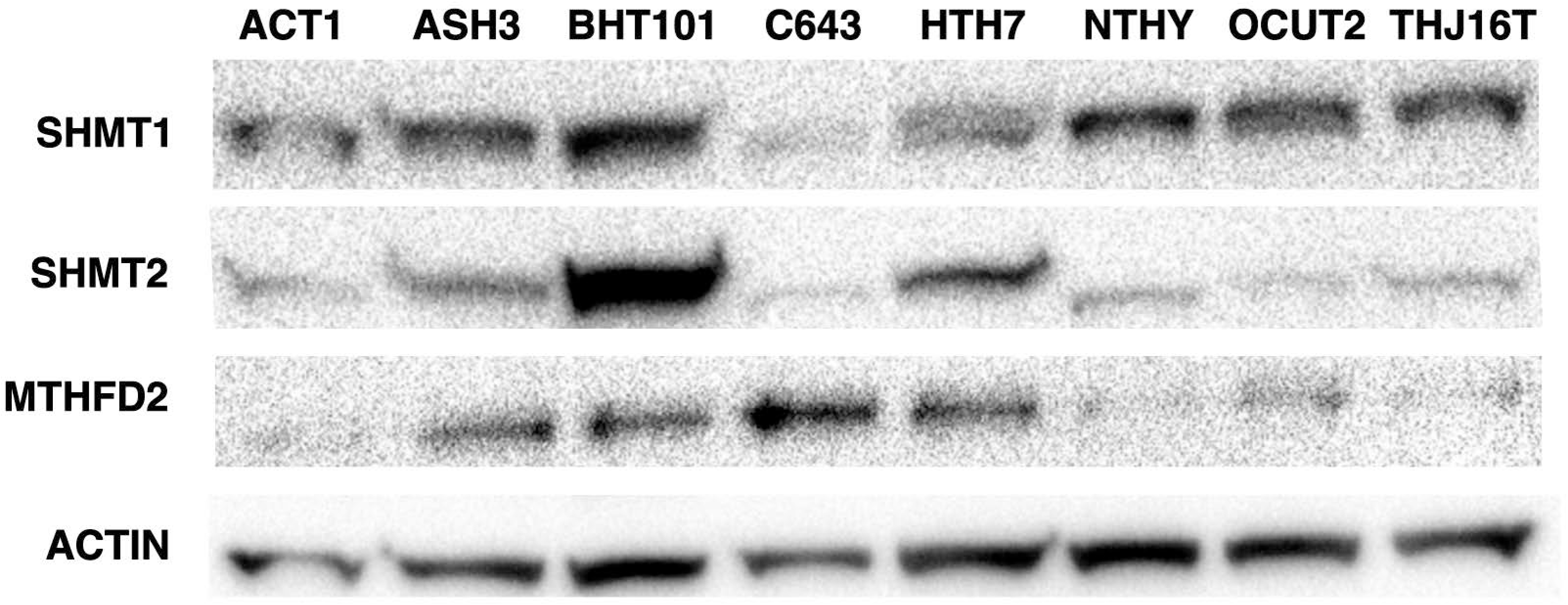
Western blot analysis of the expression of SHMT1, SHMT2, and MTHFD2 in a panel of human ATC cells lines. Immortalized NTHY-ori cells are included as a comparison for non-transformed cells.

**Suppl. Fig. 2.**
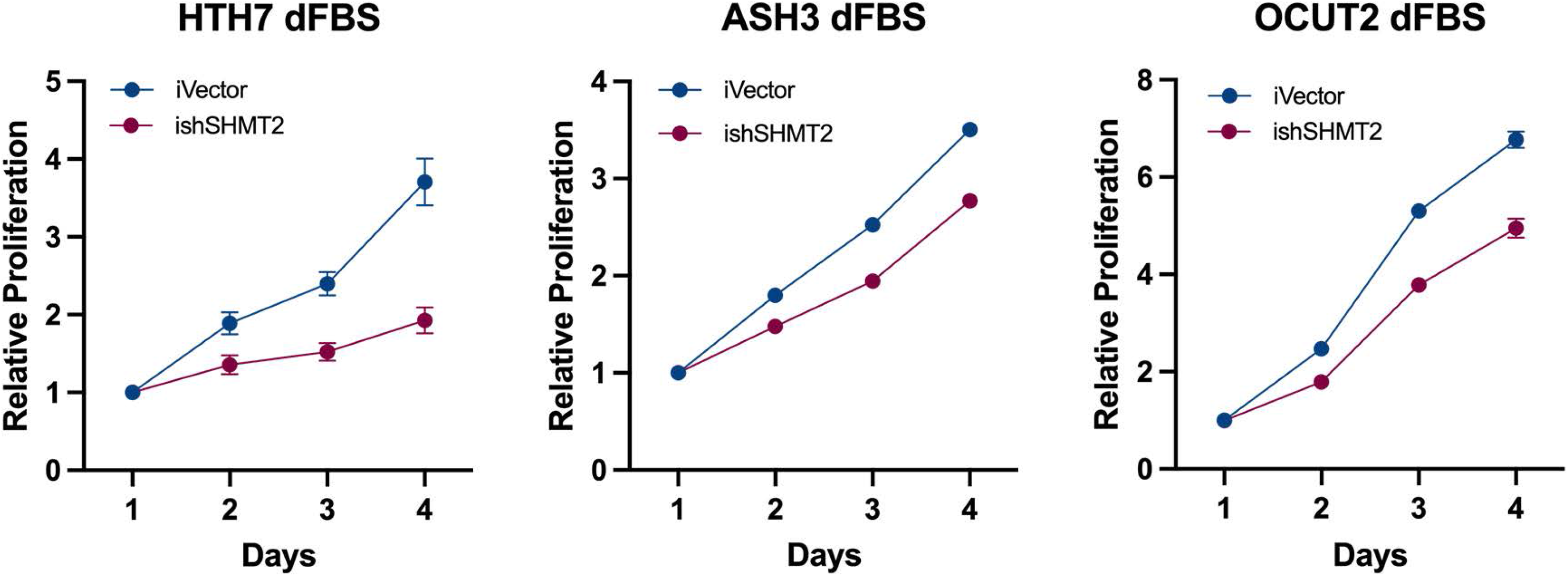
Analysis of cell proliferation in medium with dialyzed FBS of additional human ATC cell lines transduced with doxycycline-inducible non-targeting shRNA or sh*SHMT2*. Similar results were obtained with a second set of shRNAs (not shown).

**Suppl. Fig. 3.**
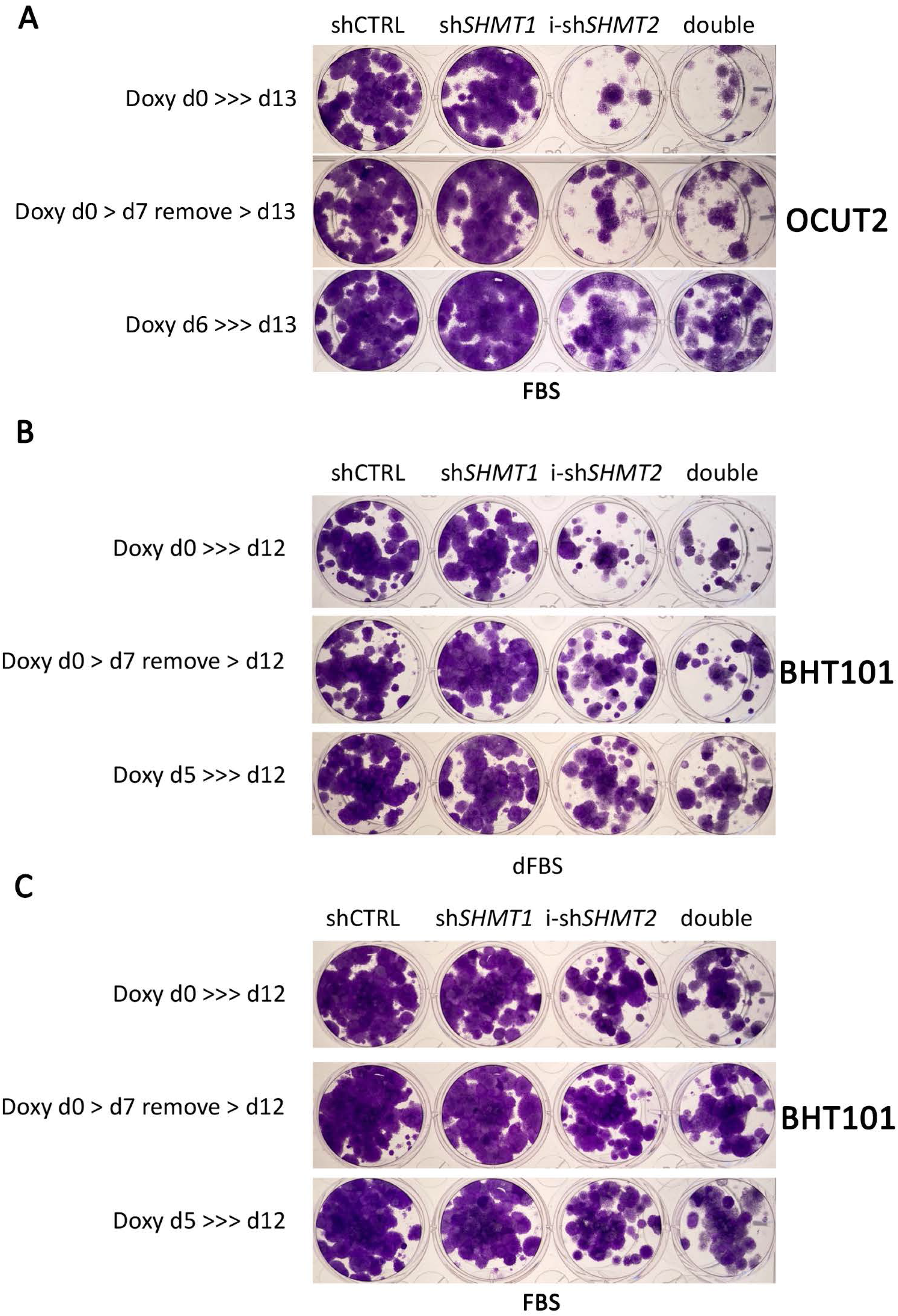
Colony forming assays of OCUT2 and BHT101 ATC cell lines expressing the indicated shRNAs. Expression of sh*SHMT2* was induced at day 0 (top and middle) or at day 5/6. In the middle panel, doxycycline was removed at day 7 to allow re-expression of sh*SHMT2*. Cells were grown with regular or dialyzed FBS as indicated.

**Suppl. Fig. 4.**
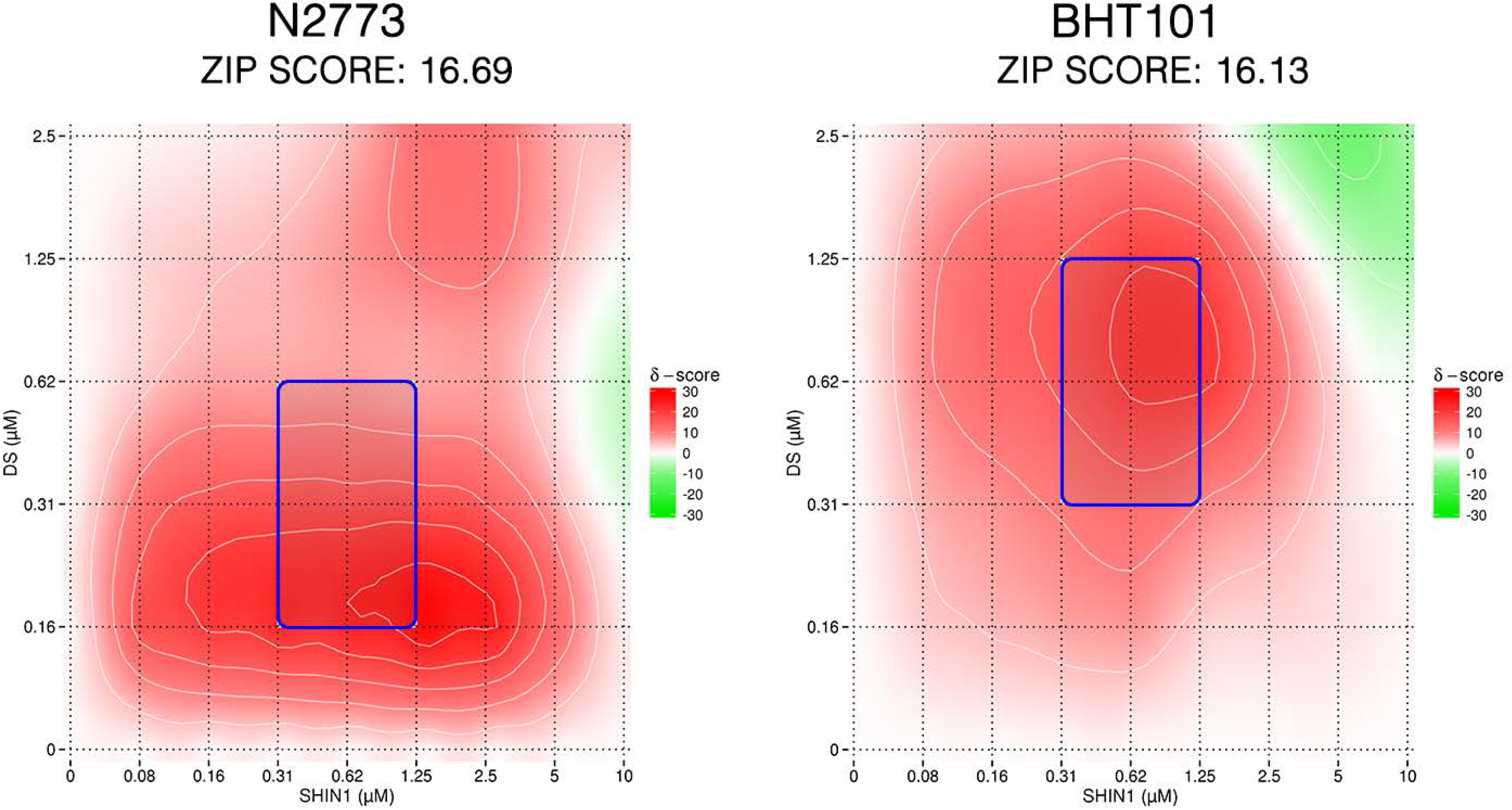
Analysis of synergy between SHMT1/2 and MTHFD2 inhibitors in mouse (N2773) and human (BHT101) ATC cell lines. Synergy was analyzed using Synergy Finder. ZIP score above 10 indicates synergy.

**Suppl. Fig. 5.**
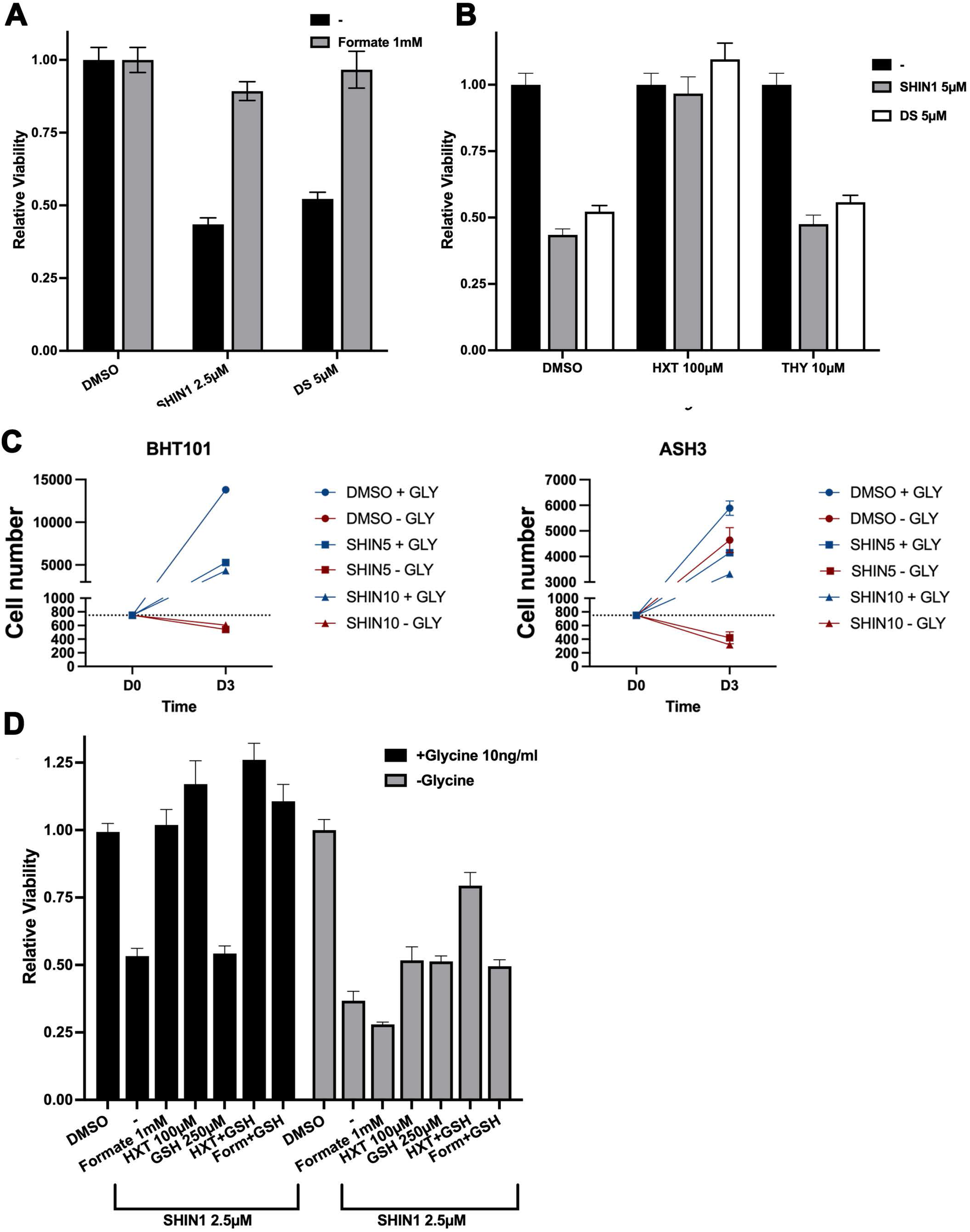
A) Rescuing effect of exogenous formate on the growth inhibition induced by one-carbon pathway inhibitors in C643 cells. B) Exogenous hypoxanthine, but not thymidine, rescues the growth inhibition induced by one-carbon pathway inhibitors in C643 cells. C) Lethal effect of exogenous glycine deprivation combined with SHIN1 in BHT101 and ASH3 cells. The dotted line indicates cell number at the time of treatment. D) Systematic analysis of the extent of viability rescue in the presence of exogenous formate, hypoxanthine, and glutathione (GSH) in the presence and absence of extracellular glycine in C643 cells.

